# Endoderm differentiates into a transient epidermis in the mouse perineum

**DOI:** 10.1101/2025.02.03.636156

**Authors:** Christine E. Larkins, Daniel M. Grunberg, Gabriel M. Daniels, Erik J. Feldtmann, Martin J. Cohn

## Abstract

In eutherian mammals, the embryonic cloaca is partitioned into genitourinary and anorectal canals by the urorectal septum. At the caudal end of the mouse embryo, the urorectal septum contributes to the perineum, which separates the anus from the external genitalia. Growth of the urorectal septum displaces cloacal endoderm to the surface of the perineum, where it is incorporated into epidermis, an enigmatic fate for endodermal cells. Here we show that endodermal cells differentiate into true epidermis in the perineum, expressing basal, spinous, and granular cell markers. Endodermal epidermis is lost through terminal differentiation and desquamation postnatally, when it is replaced by ectoderm. Live imaging and single-cell tracking reveal that ectodermal cells move at a faster velocity in a lateral-to-medial direction, converging towards the narrow band of endoderm between the anus and external genitalia. Although the perineum is sexually dimorphic, similar spatiotemporal patterns of cell movement were observed in males and females. These results demonstrate that cloacal endoderm differentiates into a non-renewing, transient epidermis at the midline of the perineum. Differential movement of endodermal and ectodermal cells suggests that perineum epidermis develops by convergent extension. These findings provide a foundation for further studies of perineum development and of sex-specific epidermal phenotypes.

## Introduction

In eutherian mammals, the openings for the digestive and genitourinary tracts are separated by the perineum, which forms the base of the pelvic floor between the anus and the external genitalia. Anogenital distance reflects the dorsal-ventral length of the perineum, which is longer in males relative to females. Anatomically, the perineum consists of deep muscular layers and a superficial layer of skin. During development, the posterior region of the embryonic hindgut, known as the cloaca, is partitioned into genitourinary and anorectal canals by the urorectal septum. At the completion of cloacal septation, when the urorectal septum reaches the caudal end of the embryo, the surface ectoderm of the cloacal membrane undergoes apoptosis and the urorectal septum fuses with the lateral cloacal swellings to form the definitive perineum (Hynes and Fraher, 2004; Nievelstein et al., 1998; Sasaki et al., 2004; Seifert et al., 2008; Wang et al., 2013).

Cell lineage analysis showed that degradation of cloacal membrane ectoderm exposes the underlying endoderm and that posterior growth of the urorectal septum drives the endodermal epithelium towards caudal surface of the embryo, where it is incorporated into the epidermis along the midline of the perineum (Seifert et al., 2008). This endodermal population of the perineum epidermis persists, at least through postnatal day (P) 0, along the epidermal seam, or raphe, which extends from the anus to the base of the vulva in females and to the frenulum of the penis in males (Seifert et al., 2008). Thus, the perineum is an enigmatic example of endoderm contributing to epidermis, as elsewhere in the embryo, the epidermis develops from surface ectoderm.

The epidermis of the skin consists of a stratified epithelium that serves as a barrier for protection from physical, chemical, thermal, and pathogenic insults, responds to environmental stimuli, prevents water loss, and regulates body temperature. The surface ectoderm covering the embryo is transformed from a single-layered (simple) epithelium to the stratified (complex) epithelium of the epidermis as cells divide perpendicular to, or detach from, the basement membrane and migrate upward from the basal layer, giving rise to the spinous, granular, and cornified layers (Lechler and Fuchs, 2005; Smart, 1970). Epidermis is a self-renewing tissue; progenitor cells originate from the basal layer, where the stem cells reside, and terminally differentiate as they move through the spinous and granular cell layers towards the surface, where they enucleate to form the stratum corneum of the epidermis (Blanpain and Fuchs, 2009; Koster and Roop, 2007). It is through formation of the stratum corneum that the skin acquires barrier properties. The superficial layers of the stratum corneum are continuously lost through desquamation and are replaced by basal cells that divide, stratify, and differentiate (Elias, 1983; Rawlings et al., 1994).

Sex differences are found in numerous organs, including the skin, where thickness, immunity, hair growth, sweat gland number, lipid content, and collagen composition differ between males and females (Arai et al., 2017; Azzi et al., 2005; Chi et al., 2024; Sandby-Moller et al., 2003; Shuster et al., 1975; Zouboulis et al., 2007). Skin diseases also show sex biases. For example, neoplasias and infectious diseases, including melanoma, actinic keratosis, and leishmaniases, generally occur more often in men, whereas autoimmune and autoinflammatory diseases, such as systemic and cutaneous lupus, dermatomyositis, and psoriasis, are more prevalent in women (Cosci et al., 2022; Dao and Kazin, 2007; de Araújo Albuquerque et al., 2021; Giacomoni et al., 2009; Li et al., 2022; Lopes Almeida Gomes et al., 2024; Zheng et al., 2022). A number of studies showed that androgens and/or estrogens can impact the risk or progression of skin diseases and play roles in sexually dimorphic development of skin (Arai et al., 2017; Azzi et al., 2005; Hanley et al., 1996). The size of the perineum (measured as anogenital distance) is sexually dimorphic due to androgen signaling, which increases the length of the perineum (Hsieh et al., 2012; Macleod et al., 2010; McDermott et al., 1978).

In this study, we examined the fate of the endodermal lineage and the cellular behaviors that underlie development and sexual differentiation of the perineum epidermis. We show that endodermal cells that contribute to the perineum skin express markers of the basal, spinous, and granular layers of the epidermis in a spatially organized pattern, confirming that the endodermal lineage differentiates into true epidermis. We find that endodermal epidermis is transient, however, and eventually is replaced by the ectodermal lineage. Live imaging of endodermal and ectodermal cell behavior during development of the perineal epidermis revealed that ectodermal cells move at a faster velocity in a lateral-to-medial direction, towards the endodermal lineage, which becomes increasingly restricted to a narrow band between the anus and external genitalia. Despite the sexually dimorphic length of the perineum, cell velocities and trajectories during did not differ between males and females.

## Results

### A transient population of endoderm cells in the perineum skin

After the embryonic cloaca has been partitioned into genitourinary and anorectal sinuses by the urorectal septum (at E13.5 in mice), the cloacal membrane ruptures, exposing the underlying endodermal epithelium at the caudal end of the urorectal septum, which forms the central margin of the perineum (Seifert et al., 2008). To determine whether the endodermal lineage of the perineal epithelium persists as a self-renewing epidermis, we used the *Shh^GFPcre^* allele to irreversibly activate the *Rosa26^lacZ^* reporter in endodermal cells, and we followed their fate through postnatal stages (Figure 1). At E14.5, endoderm cells showed similar distributions in the perineum of *Shh^GFPcre^*^/+^; *R26R^LacZ^*^/+^ males and females (Figure 1A,B). The endodermal domain then narrows and elongates along the dorsoventral axis of the perineum (between the anus and genital tubercle), with males showing a greater degree of elongation and narrowing compared to females after E14.5 (Figure 1C-F; Seifert 2008). By P3, the endodermal domain of the perineum had become discontinuous in males, whereas females showed a robust stripe of endodermal cells between the vaginal opening and the anus (Figure 1G,H).

**Figure 1.**
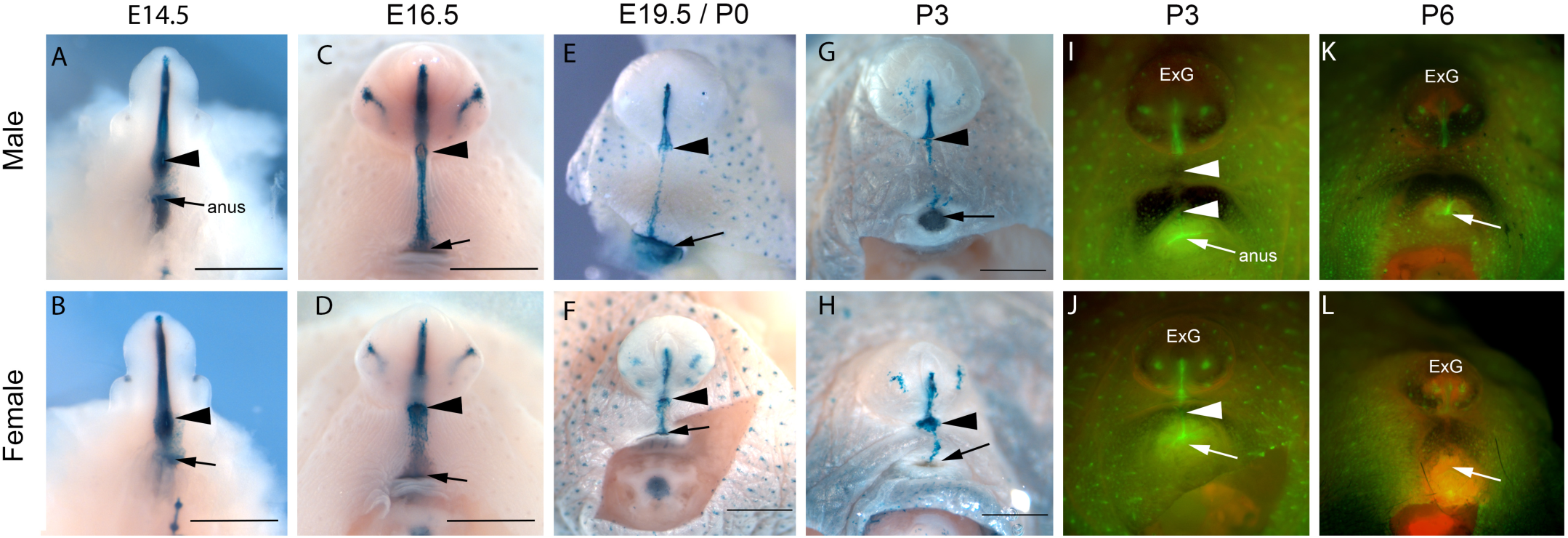
Endoderm cell lineage tracing in the perineum of male and female mice. (A-H) *Shh^GFPcre/+^; R26R^lacZ/+^* embryos and pups stained with X-gal to show *Shh*-expressing cells and their descendants in the endodermal lineage of the perineum from E14.5 to P3. Black arrowheads mark the base of the genital tubercle/phallus and black arrows indicate the position of the anus. Scale bar is 1mm. (I-L) *Shh^GFPcre/+^; R26R^mTmG/+^* males and females at P3 and P6 showing GFP expression in the *Shh*-expressing cells and their descendants. White arrowheads mark GFP-positive cells at the midline of the perineum; white arrows mark the position of the anus; ExG, external genitalia.

Given that penetration of staining solutions can be limited after the epidermal barrier has developed, we also examined the endodermal lineage at postnatal stages by conditional activation of the *Rosa26^mTmG^* fluorescent reporter, which allows visualization of membrane-bound GFP in endodermal cells following *cre* expression. In whole mount *Shh^GFPcre/+^; Rosa26^mTmG/+^* specimens, we observed a reduction in the number of GFP- positive cells in the perineum after birth, and expression became undetectable between P3 and P6 (Figure 1I-L). GFP could be seen at both stages in other *Shh*-expressing tissues, such as the urethra, hair follicles, and preputial glands. These results show that endodermal cells contribute to the midline of the perineum epidermis, but this lineage is transient and is lost during the first week of postnatal development.

### Endoderm cells in the perineum differentiate into *bona fide* epidermis

To determine if the endoderm cells in the perineum skin acquire true epidermal identity, we examined markers of differentiating epidermal cells. Between E15.5 and E17.5, epidermal differentiation markers were detected first in ectodermal cells of the lateral perineum, and expression gradually expanded medially as expression was activated in the endodermal lineage (Figures 2 and S1). In males and females at E15.5, ectodermal cells were positive for keratin 10 (K10), which marks epidermal cells in the spinous layer, and for loricrin and filaggrin, which mark the granular layer, but endoderm cells at the midline showed weak or no expression of these epidermal markers (Figures 2A-D and S1A,B). In males at E16.5, K10 expression had expanded medially and became contiguous across the ectoderm and the endoderm, which had contracted further medially (Figure 2E). Females at E16.5 also showed expression of K10 in ectodermal and endodermal cells, although the endodermal lineage was wider than in males and K10 was detected in lateral but not medial endodermal cells (Figure 2F). In E16.5 embryos of both sexes, ectoderm and lateral endoderm cells were positive for loricrin, but the medial endoderm, which was narrower in males, showed little or no signal (Figure 2G,H). Filaggrin expression remained weak in the perineum epidermis at E16.5 and, as with the other markers, a gap in expression was observed in the medial endodermal cells (Supplemental Figure S1E,F). By E17.5, activity of these spinous and granular markers had expanded further medially to form a contiguous domain of expression throughout the perineum epidermis, such that endoderm cells within these layers now expressed K10, loricrin, and filaggrin (Figure 2I-L and Supplemental Figure S1I,J). We also examined the keratinocyte marker cytokeratin 14 (K14) in developing perineum epidermis from E15.5 to E17.5 and found that both endodermal and ectodermal cells expressed K14 at all stages (Figure S1C,D, G,H, and J,L) At E15.5, K14 was detected in basal and suprabasal layers (Supplemental Figure S1C,D), but expression became restricted to the basal layer by E17.5 (Supplemental Figure S1K,L). Together, these results indicate that endodermal cells at the midline of the perineum acquire epidermal identity, although there is a delay in their differentiation relative to the adjacent ectoderm. Our data from the perineum epidermis are consistent with previous reports that epidermal differentiation occurs in waves from dorsal to ventral (Hardman et al., 1998)

**Figure 2.**
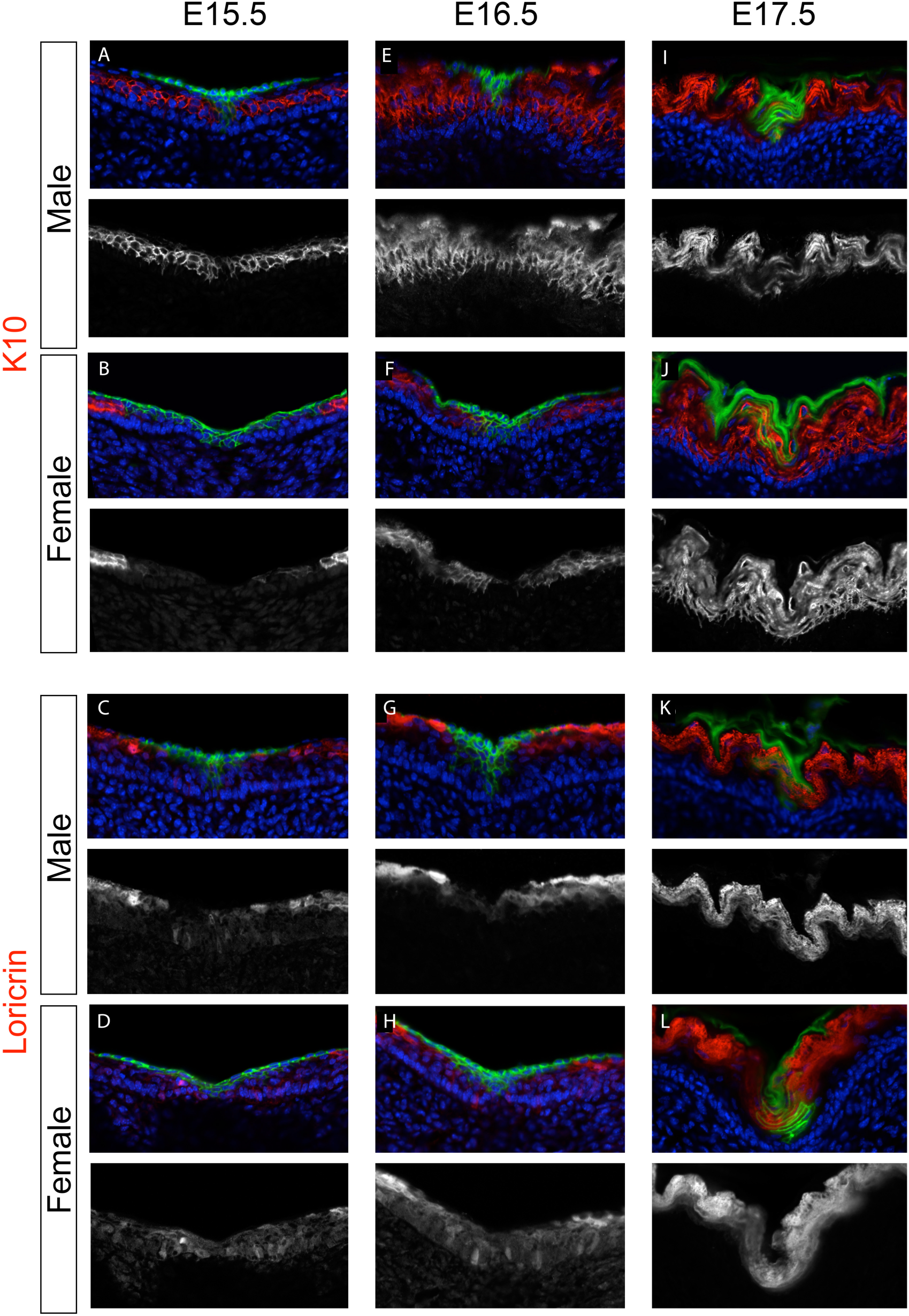
The endodermal lineage of the perineum expresses epidermal differentiation markers. (A-L) *Shh^GFPcre/+^*; *R26R^mTmG/+^* males and females at E15.5, E16.5, and E17.5 showing immunofluorescence of K10 (A,B,E,F,I,J) and loricrin (C,D,G,H,K,L). Green signal indicates GFP expression in endoderm. Red signal is pseudocolored immunofluorescence of K10 and loricrin. Blue indicates Hoechst staining of cell nuclei. Greyscale panels show single-channel immunofluorescence.

### Terminal differentiation leads to postnatal loss of the endodermal lineage from the perineum epidermis

Epidermis consists of distinct layers of cells that undergo terminal differentiation and are continuously replaced by a self-renewing population of epidermal stem cells in the basal layer. Our finding that cloacal endoderm gives rise to a population of epidermal cells in the perineum that is lost between P3 and P6 suggested that the endodermal lineage undergoes terminal differentiation and desquamation and is replaced by a different lineage of cells. To test this hypothesis, we traced the fate of the endodermal population of the epidermis from E14.5 through P6 in sections through the perineum of *Shh^GFPcre^*; *R26R^mTmG/+^* males and females (Figure 3). At E14.5, when the perineum epidermis is 2-3 cell layers thick, GFP+ endoderm cells were found in basal and suprabasal layers at the midline of the perineum and in a broader domain of suprabasal layers lateral to the midline (Figure 3A,B). By E16.5, GFP+ endoderm cells were detected in all suprabasal layers of the perineum epidermis at the midline, but very few GFP+ cells were attached to the basement membrane (Figure 3C,D). In more lateral regions of the perineum, endodermal cells were found only in the stratum corneum, the most superficial layer of the epidermis (Figure 4C,D). When we examined the epidermis of perinatal (E19.5-P0) and early postnatal (P3) males and females, the endodermal lineage was found predominantly in the stratum corneum (Figure 3E-H). By P3, the endodermal domain had contracted medially and was restricted to a narrow domain of superficial cells at the midline of the perineum (Figure 3G,H). The basal, spinous, and granular layers were almost completely devoid of GFP+ cells at P3; a few isolated endoderm cells were detected in some sections, but the majority of endodermal descendants were confined to the stratum corneum (Figure 3G,H). By P6, the stratum corneum was negative for GFP, indicating that the endodermal lineage had been sloughed (Figure 3I,J). At this stage, GFP expression could be seen only in developing hair follicles, which are known to express *Shh* (St-Jacques et al., 1998). Thus, the endodermal population of the perineum epidermis is transient rather than self-renewing. These results suggest that endodermal cells undergo terminal differentiation and desquamation in a lateral-to-medial direction and are eliminated from the epidermis of the perineum by P6.

**Figure 3.**
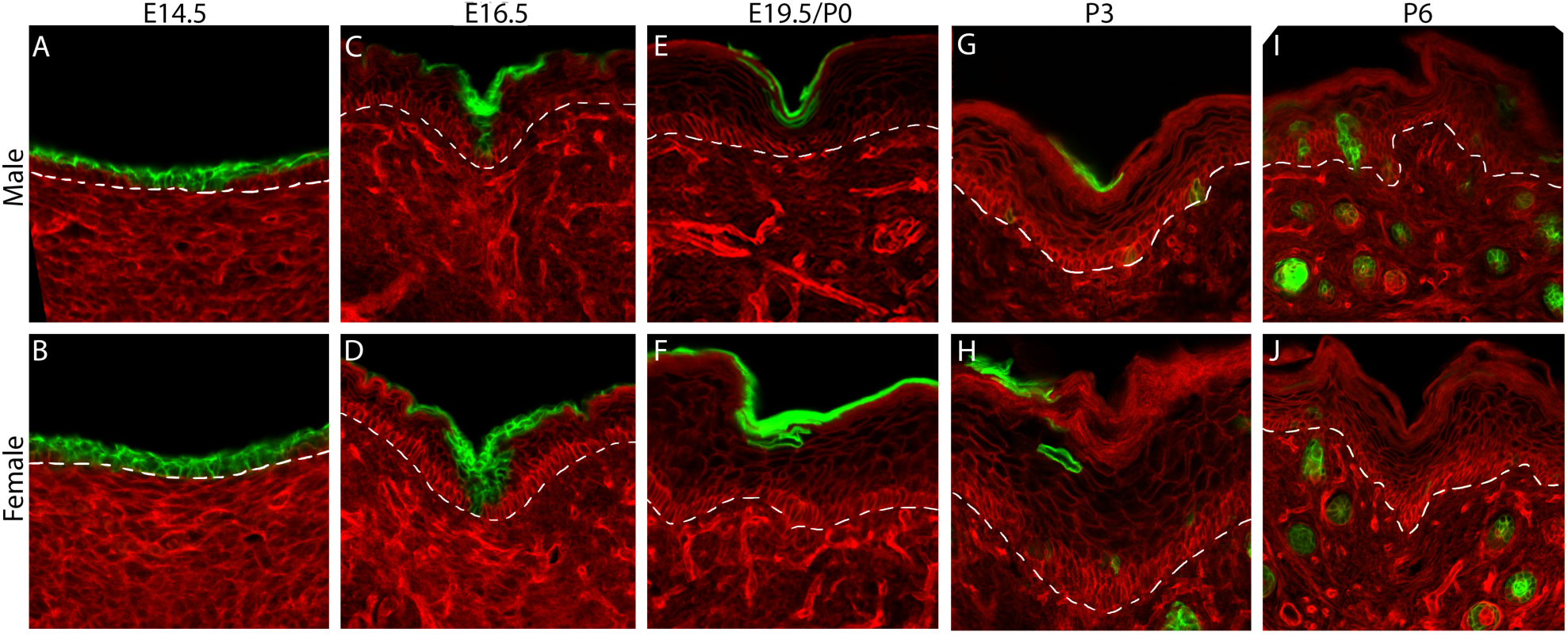
Terminal differentiation leads to loss of endoderm from the epidermis. (A-J) Transverse sections through the perineal skin of male and female *Shh^GFPcre/+^*; *R26R^mTmG/+^* mice from E14.5-P6. (A,B) At E14.5, GFP-expressing endodermal cells are found in the basal and suprabasal layers of epidermis in the medial region of the perineum. (C,D) At E16.5, GFP-positive endoderm is throughout all epidermal layers at the midline, but laterally, the majority of endodermal cells are located suprabasally. (E,F) At E19.5-P0, the endoderm lineage is almost entirely in the stratum corneum and no GFP-positive cells are detectable in the basal layers. (G-J) At P6, a small domain of endodermal cells is visible in the stratum corneum at the midline of the perineum epidermis (G,H), but these are no longer detectable at P6 (I,J). GFP expression in the in the hair follicles is seen in the dermis.

**Figure 4.**
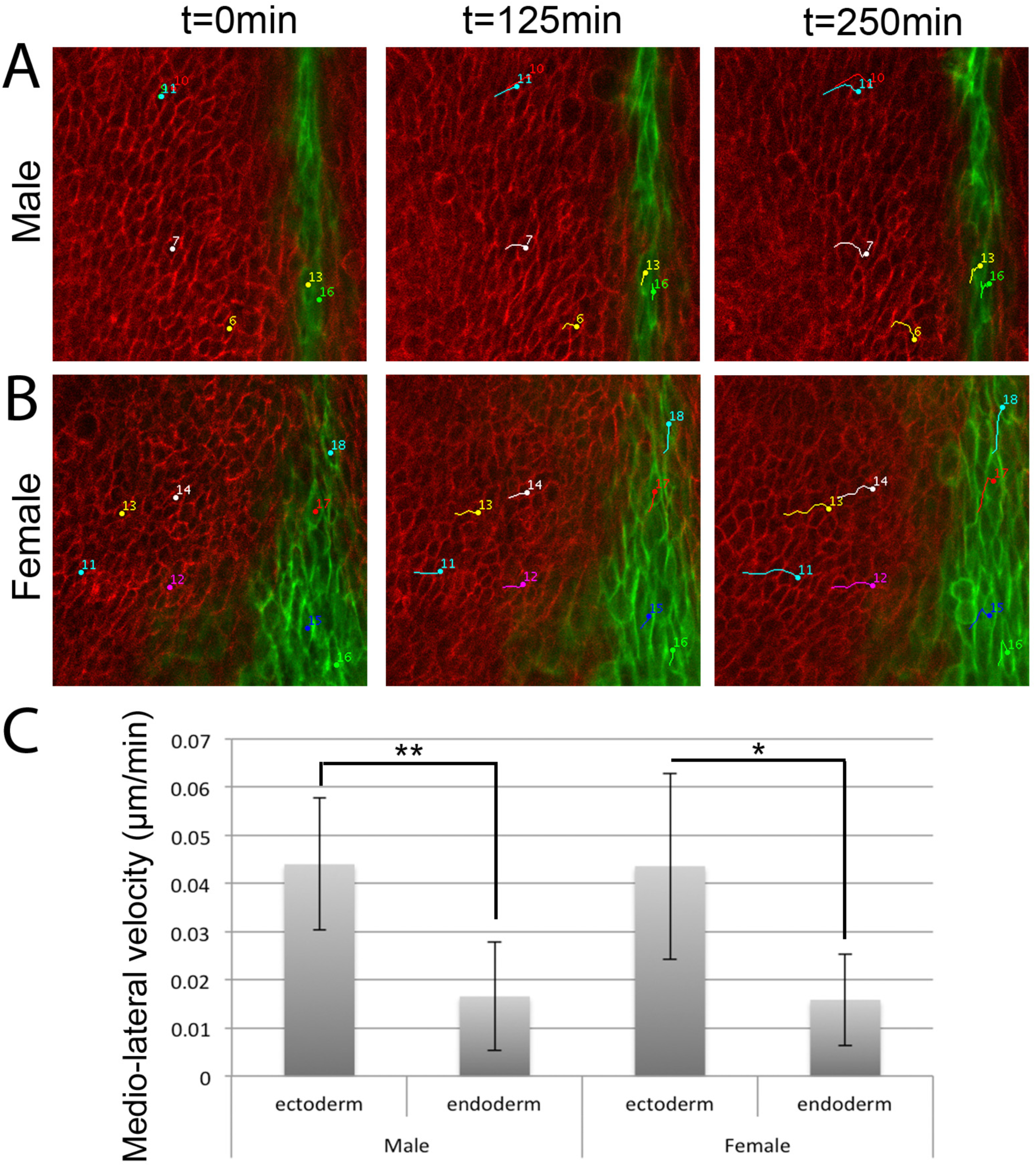
Live imaging of cell movements in the endoderm and surrounding ectoderm of the perineum. (A,B) Time lapse of a male and a female *Shh^GFPcre/+^*; *R26R^mTmG/+^* perineum. Cell tracks are marked with colored dots and lines. Cells were followed on both sides of the midline of the perineum, although only one side is shown here. (C) Graph showing the average mediolateral velocity of ectoderm and endoderm cells. Note that ectoderm cells have a significantly higher mediolateral velocity compared to endoderm cells \**P* < 0.05 (*P* = 0.015); \*\**P* < 0.01 (*P* = 0.003), using a two-tailed t test.

### Lineage-specific differences in cell movement show convergent extension of the perineum epidermis

The sexually dimorphic size of the perineum is evident in the length of the anogenital distance, but the cellular processes that drive morphogenesis and sexual differentiation of the perineum are not well understood. In males, the endodermal domain of the perineum epidermis is longer (dorsoventrally) and narrower (mediolaterally) compared to females (Figure 1C-H). This observation led us to test two hypotheses; (1) that the perineum epidermis undergoes convergent extension, and (2) that patterns of epidermal cell movement differ between males and females. We performed live-imaging and single-cell tracking of endoderm and ectoderm cells in males and females at E15.5, when endoderm cells are detectable along the midline of the perineum (Seifert et al., 2008). Analysis of epidermal cell trajectories and velocities in the developing perineum of males showed that ectoderm cells move medially, towards the endoderm cells, at a rate of 0.044 μm/min (Figure 5 A, C). Endoderm cells, by comparison, showed significantly slower mediolateral movement (0.017 μm/min) when compared to ectodermal cells (*P*=0.004; Figure 5A,C). However, when we compared the total velocity of cell movement in any direction, ectoderm and endoderm cells were not significantly different (0.089 and 0.086 um/min, respectively; *P*=0.7). Thus, ectoderm and endoderm cells of the perineum epidermis moved the same total distance, but ectoderm cells preferentially migrate along a lateral-to-medial trajectory, converging towards endoderm cells at the midline of the perineum. We then conducted an identical analysis of E15.5 females to determine whether sexual dimorphism of the anogenital distance is associated with sex differences in cell movement during development of the perineum epidermis. The rates of mediolateral movement by ectodermal and endodermal cells in the female perineum epidermis were identical to those observed in males (Figure 5 B,C). Taken together, these results show that morphogenesis of the male and female perineum epidermis involves convergence of the lateral ectodermal lineage towards the medial endodermal lineage.

## Discussion

The three germ layers of the embryo give rise to distinctive tissues. Generally, ectoderm gives rise to epidermis of the skin and neural derivatives, and endoderm forms the epithelial lining of visceral organs. In previous studies, we showed that endoderm cells at the caudal end of the hindgut are incorporated into the skin of the perineum, which is an enigmatic fate for endodermal cells. The results presented here show that this population of endoderm cells differentiates into *bona fide* epidermis that expresses the same molecular markers that characterize ectoderm-derived epidermis. We also find that although the endoderm gives rise to epidermis at the midline of the perineum, endodermal epidermis is not self-renewing. Rather, endodermal cells of the perineum epidermis undergo terminal differentiation and are lost during the neonatal period, when they are replaced by adjacent ectoderm cells.

Given that gut endoderm is an epithelium that responds to mesenchymal cues to determine its identity, it is not surprising that the population of cloacal endoderm that is displaced to the posterior surface of the embryo would acquire an epidermal fate in the context of surrounding ectoderm and underlying dermal mesenchyme (Jerman et al., 2015; Spence et al., 2011; Thomson et al., 2002; Wells and Melton, 2000). However, because the endoderm lineage is lost from the epidermis at postnatal stages, it likely lacks the stem-ness required to regenerate and maintain itself as epidermis in the perineum. There has been debate regarding the origin of the stem cells that regenerate the interfollicular epidermis and there appear to be differences in the regenerative potential of this tissue in its embryonic versus adult states (Levy et al., 2005; Sada et al., 2016; Schepeler et al., 2014; Snippert et al., 2010). Some studies suggested that there are slow cycling cells within the interfollicular epidermis, while others argued that any cell in the basal layer has the capacity to self-renew indefinitely (Clayton et al., 2007; Mascre et al., 2012; Rompolas et al., 2016). Our results show that the endoderm-derived basal epidermal cells at the midline of the perineum are unable to self-renew indefinitely. Why the endodermal lineage of the perineum epidermis lacks the stem cell properties found in ectodermal epidermis requires further investigation.

Our live imaging and cell tracking of ectodermal and endodermal cells during development of the perineum epidermis showed that ectoderm cells move along a medial trajectory, converging towards the endoderm cells at the midline. Endoderm cells move at the same velocity as ectoderm cells, but the former lack the polarized directionality of the latter. These patterns of cell movement raise the possibility that morphogenesis of the perineum epidermis involves convergent extension, in which ectodermal cells of the epidermis converge medially towards the endodermal epidermal cells at the midline, which becomes compressed into the narrow band of cells that extends between the anus and external genitalia. We tested for sex differences in these morphogenetic movements because the perineum is sexually dimorphic, with males having a longer perineum along the dorsoventral axis. Based on previous reports that the endodermal domain of the perineum epidermis becomes longer and narrower in males relative to females (Seifert et al., 2008), and that a fusion event occurs during development of the perineum in males but not in females (Glenister, 1954; Jin et al., 2016), we hypothesized that sex differences in epithelial convergent extension contribute to its dimorphic size. However, live imaging of the endodermal and ectodermal lineages of the perineum showed no difference in cell movements between males and females. Our results suggest that convergent extension of the epidermis is an integral part of perineum development in both sexes and does not account for its dimorphic length. While we cannot exclude the possibility that the apparent lack of sex differences in epidermal cell movement reflects limitations in our cell imaging and/or tracking methods, we suggest that sexual dimorphism of the perineum is likely to be a consequence of differential development of the pelvic floor rather than the epidermis.

Another possible explanation for the sexually dimorphic size of the endodermal domain of the perineum is that the epidermis differentiates more rapidly in males than in females. Previous studies comparing barrier formation and epidermal thickness between sexes reported that females show more rapid formation of the epidermal barrier and have thicker epidermis, but, to our knowledge, the rate of cell turnover of the epidermis has not been compared between sexes (Azzi et al., 2005; Hanley et al., 1996; Moverare et al., 2002). It is possible that the epidermis of the female perineum has slower cell turnover, and, as a result, perineum epidermal cells are retained for longer in females relative to males. Future experiments examining rates of endodermal and ectodermal cell differentiation and turnover in the perineum epidermis of both sexes are needed to tease apart these possibilities.

## Methods

### Mice

The *Shh^tm1(EGFP/cre)Cjt^* mouse line (abbreviated *Shh^GFPcre^*) was provided by Brian Harfe, University of Florida, Gainesville, FL. The *Gt(Rosa)26^tm1Sor^* (abbreviated *R26R^lacZ^*) and *Gt(Rosa)26^tm4(ACTB-tdTomato,-EGFP)/Luo^* (abbreviated *R26R^mTmG^*) were purchased from the Jackson Laboratory (Harfe et al., 2004; Muzumdar et al., 2007; Soriano, 1999).

### LacZ staining

Embryos were fixed for one hour in 4% paraformaldehyde (PFA) at room temperature and then washed three times for twenty minutes each in lacZ buffer (0.1M phosphate buffer pH 7.4, 1% sodium deoxycholate, 2 mM MgCl_2_, 0.2% IGEPAL CA-630). The tissue was incubated overnight at room temperature in *lacZ* staining solution (lacZ buffer with X-gal, K_3_Fe(CN)_6_, and K_4_Fe(CN)_6_). Finally, phosphate buffered saline (PBS) was used to stop the reaction and the samples were fixed overnight in 4% PFA.

### Immunofluorescence

For immunofluorescence, embryos were fixed in 4% PFA for 1 hour and then washed in PBS for 2 hours. Embryos were washed 3 times for 10 minutes and embedded in OCT, then the blocks were frozen and cryosectioned (20 μm). In preparation for immunofluorescence, sections were fixed in 4% PFA for 10 minutes and then washed in PBS conaining 1% goat serum and 0.1% triton X-100 solution (wash buffer) prior to primary antibody application. Sections were incubated with primary antibodies K10 (rabbit, 1:1000, Biolegends), loricrin (rabbit, 1:750, Biolegends), and filaggrin (rabbit, 1:1000, Biolegends) at 4°C overnight in a humid box. After 3 washes in wash buffer, slides were incubated with secondary antibodies coupled to Alexa Flour 647 (Invitrogen) at 1:300 with Hoechst (1:3000) for 1 hour at room temperature and then washed in wash buffer. Sections were cover slipped with Fluoromount G and imaged on a Zeiss LSM 710 confocal microscope.

### Live Imaging

For live imaging experiments, the perineum with the genital tubercle was removed from the embryo. Organ culture was performed according to published methods (Mort et al., 2010). Briefly, the tissue was placed in a 35mm lumox membrane dish (Sarstedt), such that the perineum epidermis was in contact with the membrane. The tissue was anchored by covering it with a Nuclepore membrane (Fisher, 09-300-57) that was then covered with 500ul Matrigel that had been thawed on ice (Corning 356231, phenol red free, growth factor reduced). After incubating at 37°C for 30 minutes to allow the matrigel to solidify, the dish was filled with 1.5mL culture media [DMEM/F12 (phenol red free), 10% charcoal dextran stripped fetal bovine serum, 1% ITS (insulin, transferrin, selenium), 1x penicillin/streptomycin]. Male cultures included 10nM final concentration DHT. The culture dish was placed on a heated microscopy stage that was surrounded by a humidified 37°C chamber with 5% CO_2_ and was allowed to rest for 30 minutes prior to imaging. Z-stack images were acquired every 25 minutes on a Zeiss LSM 710 confocal microscope.

### Cell tracking analysis

LSM files were imported into ImageJ (Fiji). After merging the channels and stacking to RGB, Z-slices from the stack were selected to ensure that the basal layer of epidermal cells could be seen in 10 frames. A single Z-plane was used for cell tracking. Tracking was done using the Manual Tracking plugin, and only cells that could be followed for 10 frames were measured. Manual tracking showed the number of pixels that a cell moved in both the X and Y planes over time. This was used to calculate the velocity in the medio-lateral and dorsoventral directions. Significance was determined by performing a two-tailed t-test comparing average velocity of the endoderm and ectoderm from 6 males and 6 females.

## Acknowledgements

We thank Emily Merton, Ana Enriquez, and Brooke Armfield for assistance, Ashley Seifert for contributions to our understanding of perineum development, Brian Harfe for providing the *Shh^GfpCre^* mouse line, and the University Scholars Program at the University of Florida for supporting undergraduate student participation in this research. This work was supported by the NIH/National Institute of Diabetes and Digestive and Kidney Diseases grant number K01DK105077 to C.E.L. and R01DK110408 and U01DK131548 to M.J.C.

## Author Contributions

C.E.L. designed the experiments, performed the cell lineage, live imaging, and immunofluorescence studies, analyzed and interpreted the data, and drafted the paper. D.M.G., G.M.D., and E.J.F. contributed histological and immunofluorescence data. M.J.C. contributed to data interpretation and editing of the paper.

---

**Supplemental Figure S1.**
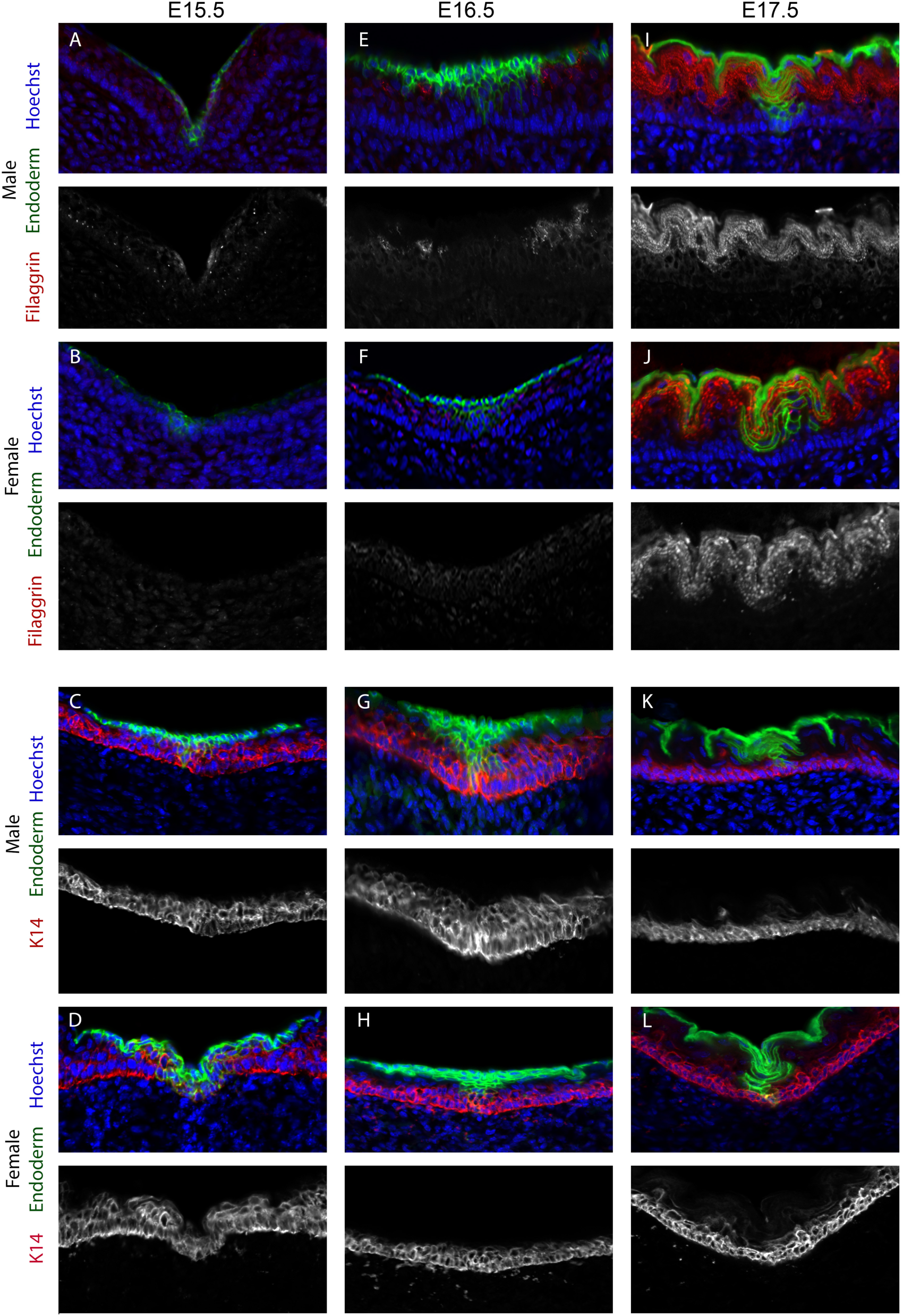
Endodermal epidermis of the perineum expresses filaggrin and K14. (A-J) *Shh^GFPcre/+^*; *R26R^mTmG/+^* males and females at E15.5, E16.5, and E17.5 showing immunofluorescence of Filaggrin (A,B E,F,I,J) and K14 (C,D,G,H,K,L). Green signal indicates GFP expression in endoderm. Red signal is pseudocolored immunofluorescence of filaggrin and K14. Blue indicates Hoechst staining of cell nuclei. Greyscale panels show single-channel immunofluorescence.

